# Lung-innervating neurons expressing *Tmc3* can induce broncho-constriction and dilation with direct consequences for the respiratory cycle

**DOI:** 10.1101/2023.08.24.554567

**Authors:** Jens Kortmann, Kevin Huang, Ming-Chi Tsai, Kai Barck, Amanda Jacobson, Cary D Austin, Debra Dunlap, Cecile Chalouni, Surinder Jeet, Alessia Balestrini, Elaine Storm, Mark S Wilson, Lunbin Deng, Michelle Dourado, David Hackos, Lorena Riol-Blanco, Joshua S. Kaminker, Shannon J. Turley

**Affiliations:** Genentech, South San Francisco, CA, USA

**Author notes:** Contributed equally to this manuscript (Jens Kortmann, Kevin Huang).

## Abstract

Sensory neurons of the vagal ganglia (VG) innervate lungs and play a critical role in maintaining airway homeostasis. However, the specific VG neurons that innervate lungs, and the mechanisms by which these neurons sense and respond to airway insults, are not well understood. Here, we identify a subpopulation of lung-innervating VG neurons defined by their expression of *Tmc3*. Single cell transcriptomics illuminated several subpopulations of *Tmc3+* sensory neurons, revealing distinct *Piezo2*- and *Trpv1*-expressing subclusters. Furthermore, *Tmc3* deficiency in VG neurons leads to global and subcluster specific transcriptional changes related to metabolic and ion channel function. Importantly, we show that broncho-constriction and dilation can be modulated through inhibition or activation of *Tmc3+* VG neurons resulting in a decrease or increase of end-expiratory lung volume, respectively. Together, our data show that *Tmc3* is a marker of lung-innervating neurons and may play a pivotal role in maintaining fundamental inspiratory and expiratory processes.

**Significance:** Harnessing the neuronal mechanisms that regulate lung function offers potential alternatives to existing corticosteroid treatment regimens for respiratory illness associated with acute bronchoconstriction including asthma, COPD, and emphysema. Our findings define *Transmembrane channel-like 3*, *Tmc3*, as a marker of lung-innervating sensory neurons, identify distinct subpopulations of *Tmc3*+ neurons with unique transcriptional profiles, and show that activation or inhibition of these neurons has a significant impact on airway function. Our work highlights potential avenues of novel targeted intervention in respiratory conditions driven by dysfunctional neuronal reflexes.

## Introduction

The integrity of vital organs requires simultaneously operating tissue-specific physiological responses while maintaining a high level of adaptability to potential perturbations of their steady state. Viscero-sensory afferents play a key role monitoring organ integrity by sensing the tissue environment and conveying the status of visceral organs to second order neurons located in the brainstem, a process that has been coined “interoception” (1–3). There is increasing evidence that interoceptive sensory afferents are highly heterogeneous, including dozens of intermingled sensory neuron subtypes organized in at least 37 clusters (4–6). However, while there are some well-studied markers that define neuron subsets, the specific functions of these subsets are less well understood.

As one of the largest barrier tissues in the human body, the lungs have respiratory performance under neuronal control with a higher complexity of vagal afferent innervation compared to other visceral organs (1–6). Vagal sensory afferents detect insults to airway integrity, resulting in engagement of various neuronal circuits that trigger protective reflexes like cough, swallow, vocal cord adduction, and laryngeal closure (5). Sensory afferents also detect inhaled pathogens or noxious compounds with a direct impact on airway immune responses and hyperreactivity (7–10). Some populations of sensory neurons have been well-characterized for their role in airway function. For example, a subset of neurons expressing *Piezo2* has been implicated in sensing airway stretch (11). However, the specific markers, innervation patterns, and functions for the vast majority of lung innervating sensory neurons are not well understood. Identifying and defining the functional role of different populations of sensory neurons relevant for airway broncho-constriction and dilation would provide a deeper understanding of lung function and could also provide novel biological and therapeutic hypotheses.

The advent of single cell and single nuclei RNA-sequencing technologies has enabled researchers to characterize the diversity and function of neurons found within the VG (12–15). For example, early single cell profiling work characterized the nodose ganglion neurons into two broad classes, namely LTMRs labeled by *Piezo2, Ntrk3, and P2ry1*, and polymodal nociceptors labeled by *Trpv1, Gfra1, Trpa1,* and *Npy2r* (*15*). In addition, gastrointestinal innervating *Oxtr^+^* and *Glp1r^+^*VG neurons have been shown to regulate food intake, presumably through distention signaling (14). Furthermore, a small subset of VG neurons expressing *P2ry1* mediates hallmarks of airway defense including pharyngeal swallow, apnea, expiratory reflexes, and vocal fold adduction (13). More recently, retrograde barcoding identified a number of visceral organs directly innervated by nodose neurons (12). Overall, these studies have yielded important insights into VG biology. An integrative approach to validate and extend these findings could provide deeper insights into sensory neuron function in the airways.

In this study, we found that vagal ganglia are enriched in *Tmc3* expression compared to other ganglia. Neuronal tracing further revealed *Tmc3* expression predominantly in lung-innervating neurons. Integrative analysis of single cell transcriptomic data from VG neurons identified distinct subpopulations of neurons expressing *Tmc3*, including two broad classes defined by expression of *Piezo2* and *Trpv1*. *Tmc3* deficiency in VG neurons leads to global and subcluster specific transcriptional changes involved in metabolic and ion channel function. Importantly, we show that broncho-constriction and dilation can be modulated through inhibition or activation of *Tmc3+* VG neurons, directly altering the volume of remaining air in the lungs at the end of each respiratory cycle. Together, our analysis provides insights into a specific population of lung innervating neurons and provides additional opportunities to understand, monitor and potentially manipulate lung-function in health and disease.

## Results

### *Tmc3* is specifically expressed by lung-innervating Vagal Ganglia (VG) neurons

Considering that the vast majority of lung afferents are vagal in origin (1, 4–6), we conducted bulk RNAseq analysis of VG neurons and thoracic DRG neurons with the aim of identifying VG-specific markers of lung-innervating neurons (Fig. 1A). Direct comparison of VG and T1-4 DRG neuronal transcripts revealed both novel and previously known DRG- and VG-specific markers, including *Prdm12* and *P2rx2*, respectively. *Phox2b* was highly expressed in the VG as expected (13). However, we identified Transmembrane channel-like 3 (Tmc3) as an uncharacterized gene that was also among the most robustly expressed and clearly specific to VG. This is consistent with a previous report that also identified *Tmc3* messenger as highly expressed in the jugular-nodose complex (JNC) (15). We next assessed the expression of *Tmc3* by targeted qRT-PCR of VG and 16 other selected tissues, including lungs (Fig. S1A). Interestingly, we found *Tmc3* to have a fairly restricted expression pattern, with limited expression beyond VG. Last, we used in situ hybridization (ISH) to confirm *Tmc3* expression in distinct areas of VG neuron soma in both mice (Fig. 1B) and humans (Fig. S1C), but not in murine lungs (Fig. 1B).

**Figure 1.**
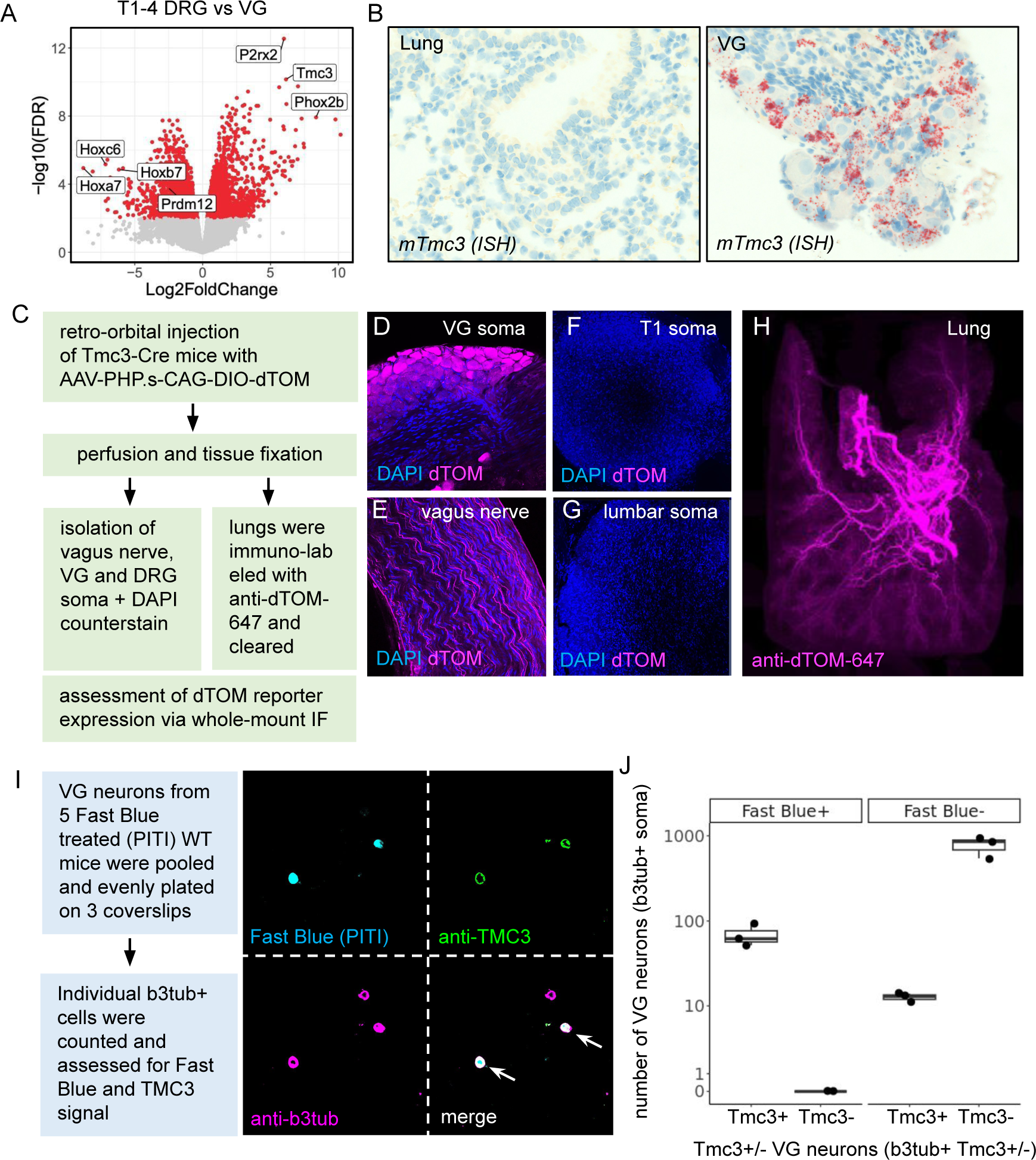
*Tmc3* is specifically expressed by lung-innervating VG neurons. (A) Bulk RNAseq highlighting differences in gene expression between thoracic DRGs and VG, including the VG-specific gene *Tmc3.* Selected nodose-specific genes indicated on plot. Red color indicates statistically significant gene expression changes (FDR < 0.01 and Fold-Change > 2). (B) Representative ISH indicating *Tmc3* transcript in sections of VG but not lung. (C) Cre-specific AAV dTomato (AAV-dT) reporter was systemically injected into adult Tmc3-Cre mice and expression of the dT reporter was detected in (D) VG soma and (E) vagus nerve, but not (F) thoracic or (G) lumbar DRG soma of Tmc3-Cre mice. (H) Innervation with dTom+ fibers observed by 3D imaging of cleared whole lungs. (I) Retrograde labeling of lung innervating neurons with PITI-delivered Fast Blue tracer (blue) and subsequent counterstain with anti-Tmc3 antibody (green) and anti-Beta Tubulin 3 (b3tub, magenta); white arrows indicate b3tub+/Tmc3+ soma. (J) Quantification of total retrograde labeled lung innervating neurons.

This prompted us to utilize an additional approach to visualize projection patterns of *Tmc3+* neurons utilizing gene specific *Cre* driver expression in combination with a novel *Cre*-dependent AAV-reporter system (16) and whole-organ three dimensional (3D) imaging. In order to achieve this, we generated Tmc3*-Cre* mice whereby the endogenous Tmc3 promoter drives Cre expression. Tmc3*-Cre* mice were systemically injected with a Peripheral Nervous System (PNS)-specific Adeno-associated virus (AAV) expressing dTomato under a Cre-dependent promoter (AAV-php.s-CAG-DIO-dTOM; schematic in Fig. 1C). Consistent with our transcriptomics data, the Tmc3-Cre driven reporter revealed robust expression of dTomato in the soma of VG neurons (Fig. 1D), but not in the soma of thoracic or lumbar DRGs (Fig. 1F and 1G). Additionally, expression of dTomato was observed in the vagus nerve (Fig. 1E). 3D imaging of murine airways revealed *Tmc3+* neuronal airway-innervation spanning from the trachea into lobes and branches, including central and peripheral compartments of the lungs (Fig. 1H). The extent of lung-innervation by *Tmc3+* VG neurons is similar to that seen in 3D images of pan-vagal sensory afferents (VG) expressing *Vglut2* (17) (Fig. S1C). Taken together, these data indicate that *Tmc3* has an expression pattern that is specific to VG, and that *Tmc3* is a marker of lung-innervating VG neurons whose projections densely innervate the lungs.

We next generated a VG neuron suspension from wild type (WT) mice that received the retrograde labeling agent Fast Blue (FB) via Passive Intra-tracheal Inhalation (PITI, see methods) to enable visualization of the soma of neurons whose axon terminals were exposed to FB in the lung (Fig. 1I). As expected, the majority of VG neuronal soma (as labeled by beta-tubulin III) were FB-, suggesting that their axons do not terminate in the lungs. These results are consistent with previously described data showing that the majority of VG neurons innervate visceral organs other than the lung (1). However, 8-10% of VG soma displayed prominent FB labeling (FB+), identifying them as lung-projecting VG neurons. Strikingly, and consistent with 3D imaging data (Fig. 1H), counterstain with a Tmc3-specific antibody (Fig. S2) revealed that 100% of FB+ lung-innervating VG neurons expressed Tmc3 protein (Tmc3+ FB+) (Figs. 1I-J). Additionally, a small number of Tmc3+ neurons were also FB-, suggesting that some Tmc3+ neurons do not innervate the lungs. These results identify Tmc3 as a highly specific marker of lung-innervating VG neurons.

### Single cell RNA-seq meta-analysis reveals conserved transcriptional modules in *Tmc3+* vagal neurons

To better characterize *Tmc3+* neurons, we re-analyzed multiple publicly available vagal ganglion scRNA-seq datasets (12, 13, 15), and performed an integrative meta-analysis. Visualization of cell clusters after data integration highlighted that all major clusters were well represented across all datasets (Fig. 2A), suggesting few study- or batch-specific clusters. To provide additional data supporting the robustness of the integration, we used an orthogonal computational approach (18) that revealed that the majority of clusters are highly reproducible (AUROC > 0.9) across datasets. Interestingly, *Tmc3* was not expressed in jugular neurons, and was only found expressed in nodose neurons. Within the population of nodose neurons, we found that the proportion of *Tmc3+* neurons is relatively consistent across datasets (on average 33% of all nodose neurons as labeled by *Phox2b*) (Figs. 2B-C), indicating a reproducible number of *Tmc3+* neurons captured by 10X scRNA sequencing from distinct laboratories.

**Figure 2.**
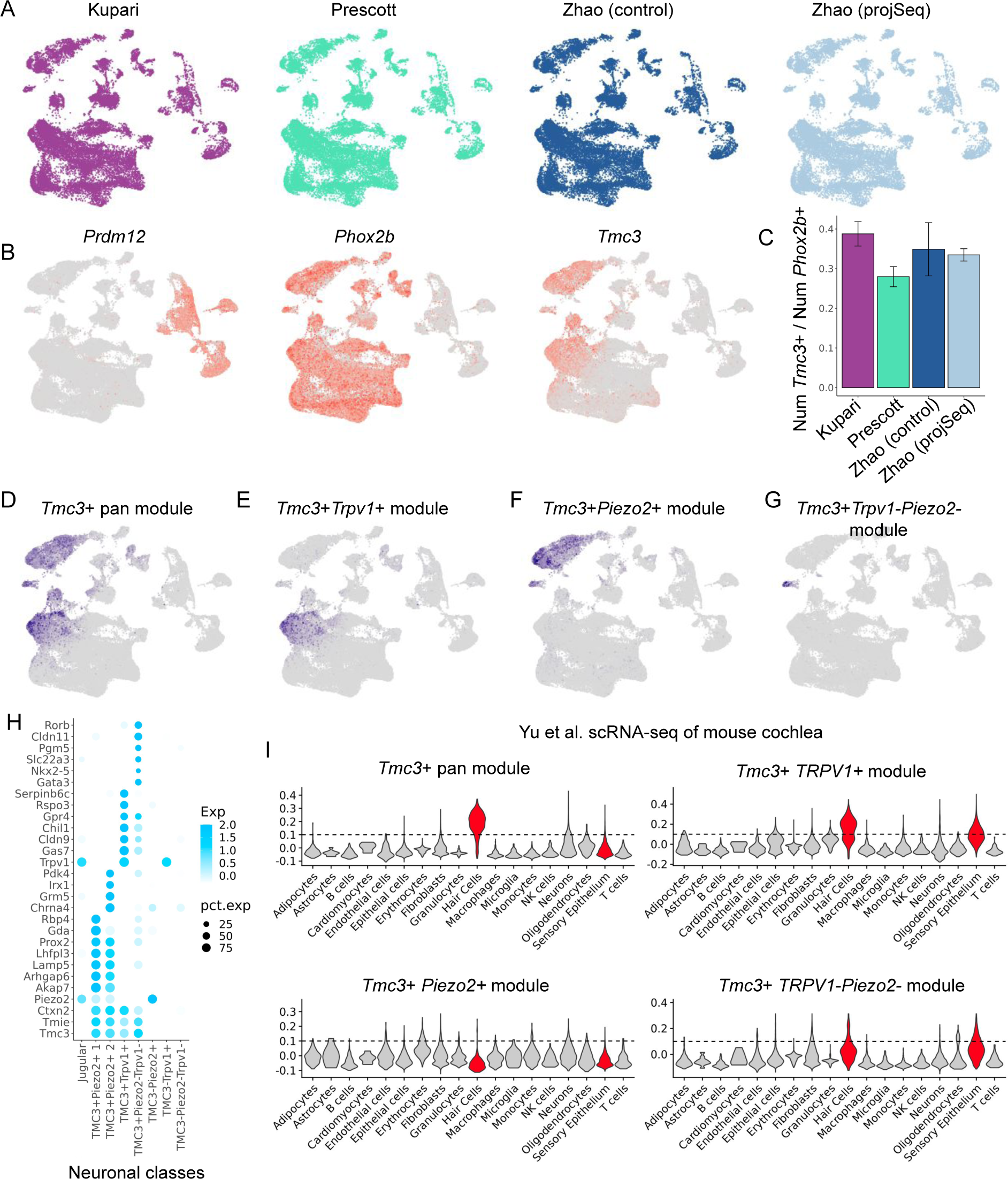
Integrative analysis of single-cell *Tmc3*+ VG neurons reveal cell population-specific gene modules. (A) UMAP representation of transcriptomic data of VG from three public datasets. Each dataset contains four biological replicate samples. (B) UMAP representation of the integrated datasets, indicating cells expressing *Prdm12*+ (jugular), *Phox2b*+ (nodose), and *Tmc3*+ neurons. Intensity of color reflects relative expression level. (C) Barplot showing the relative fraction of *Tmc3+* neurons with respect to *Phox2b*+ neurons. Error bars represent the standard deviation across 4 replicates. (D-G) UMAP representation of conserved *Tmc3*-related gene expression modules. Color reflects the intensity of gene module score. (H) Dot plot showing marker genes discriminating different subpopulations of *Tmc3*+ neurons. Size of each dot represents the percentage of each cluster expressing the respective gene (row names) and the intensity of color represents the relative expression level. (I) Violin plots of scRNA-seq showing the distribution of different *Tmc3* module scores (y-axis) across various cell populations in the inner ear (x-axis). Red color highlights cell types with elevated expression of at least one of the *Tmc3* modules.

We next performed an autocorrelation analysis (19) across all genes to identify informative gene modules that can be used to define functionally related populations of VG neurons. For this analysis, we independently identified gene modules within each dataset. We then collected those modules that were enriched for *Tmc3* expressing clusters in at least two datasets. This resulted in the identification of four gene modules with putative unique *Tmc3*-associated functionality as described below (Fig. 2D-G).

The *Tmc3+* pan gene module contains genes expressed in cells of all *Tmc3+* clusters. Notable genes in this module include *Tmie, Tmem255,* and *Ctxn2* (Fig. 2D and Fig. 2H). Interestingly, TMIE has been shown to physically interact with another TMC member, TMC1 (20), and genetic data have implicated both *Tmie* and *Tmc1* in hearing loss (20–22). The *Tmc3+Trpv1+* module contains genes expressed in a subset of the *Tmc3+* cells, and expresses *Trpv1* along with other genes including *Cldn9, Gas7,* and *Chil1* (Fig. 2E and Fig. 2H). Interestingly, mutations in *Cldn9* have been associated with hearing loss in mice and humans (23).

The *Tmc3+Piezo2+* module (Fig. 2F) expresses *Piezo2* along with other genes including *Lhflp3* and *Lamp5,* both of which have been implicated in deafness and auditory processing (24). Interestingly, we found that *Piezo2* expression was lower in *Tmc3*+ VG neurons by about 2-fold as compared to other (*Tmc3*-) *Piezo2* expressing VG neurons (Fig. 2H). Furthermore, *Tmc3+Piezo2+* neurons can further be subdivided into two subclasses; the first subclass is specifically marked by *Gda* and *Rbp4*, while the second subclass is marked by *Pdk4*, *Irx1*, *Chrna4* and *Grm5*. Finally, we identified a minor cluster of *Tmc3+* expressing cells that are neither expressing *Trpv1* or *Piezo2*. Notably, these cells express *Gata3*, which again is known to be involved in deafness. To examine potential signaling pathways that could function in *Tmc*3+ neurons, we examined neuropeptides, chemokines, and other signaling modules associated with *Tmc3*. Our analysis revealed *Galanin* as the neuropeptide with the strongest correlated with *Tmc3*+ expression in VG neurons, leading the list before other candidates like *neuropeptide B* (*NpB*), *neuropeptide Y* (*NPY*) and *vasoactive inhibitory peptide* (*VIP*) (Fig. S3A+C). Moreover, in-situ hybridization (ISH) of VG confirmed co-expression of *Tmc3* and *Galanin* (Fig. S3B).

Strikingly, the orthologs of many of the genes co-expressed with *Tmc3* have been associated with hearing loss. Due to the number of deafness genes identified in our *Tmc3* modules, we hypothesized that *Tmc3* related transcriptional programs identified in the VG may have some relationship to genes involved in auditory processing. We therefore analyzed a publicly available inner ear scRNA-seq dataset that included cells representing various stages of murine ear development (25). Intriguingly, we found that the *Tmc3*+pan and *Tmc3+Trpv1+* modules were expressed highly in inner ear hair cells (Fig. 2I). However, the *Tmc3+Piezo2+* gene signature was not found to be highly expressed in cells of the inner ear. Together, these data highlight potential conserved mechanisms between sensory neurons that innervate the airways, and the specialized cells that detect mechanosensation in the inner ear.

### *Tmc3* deficiency leads to transcriptional changes in *Tmc3*+ neuronal populations

While the function of *Tmc1* has been elucidated in a series of studies (22, 26–30), the function of the Tmc3 protein, particularly in lung-innervation, remains unknown. Based on our knowledge of lung-innervation by *Tmc3*+ VG we sought to use genetic ablation of *Tmc3* followed by NGS sequencing to elucidate the potential roles for TMC3 protein in VGs. To this end, we first performed bulk RNA sequencing from VG in WT and *Tmc3* KO mice (n=5 per group; Fig. 3A). Our analysis identified 220 and 194 genes that were up and downregulated, respectively, in *Tmc3* KO VG at homeostatic conditions (Fig. 3B and Supplementary Table 1; FDR<0.01 and LogFC > 1). Apart from *Tmc3*, *Gm16638,* a lincRNA proximal *to Tmc3,* was the second most significant RNA altered in *Tmc3* KO mice. Overall, our gene ontology analysis revealed genes downregulated in mitochondrial respiration, and upregulated in metabolic pathways (Supplementary Table 2). Of note, despite the abundance of transcriptional changes in their VG, *Tmc3* KO mice showed no obvious phenotype with respect to lung function. Hearing (auditory neural integrity) of *Tmc3* KO mice was not affected at baseline conditions (data not shown).

**Figure 3.**
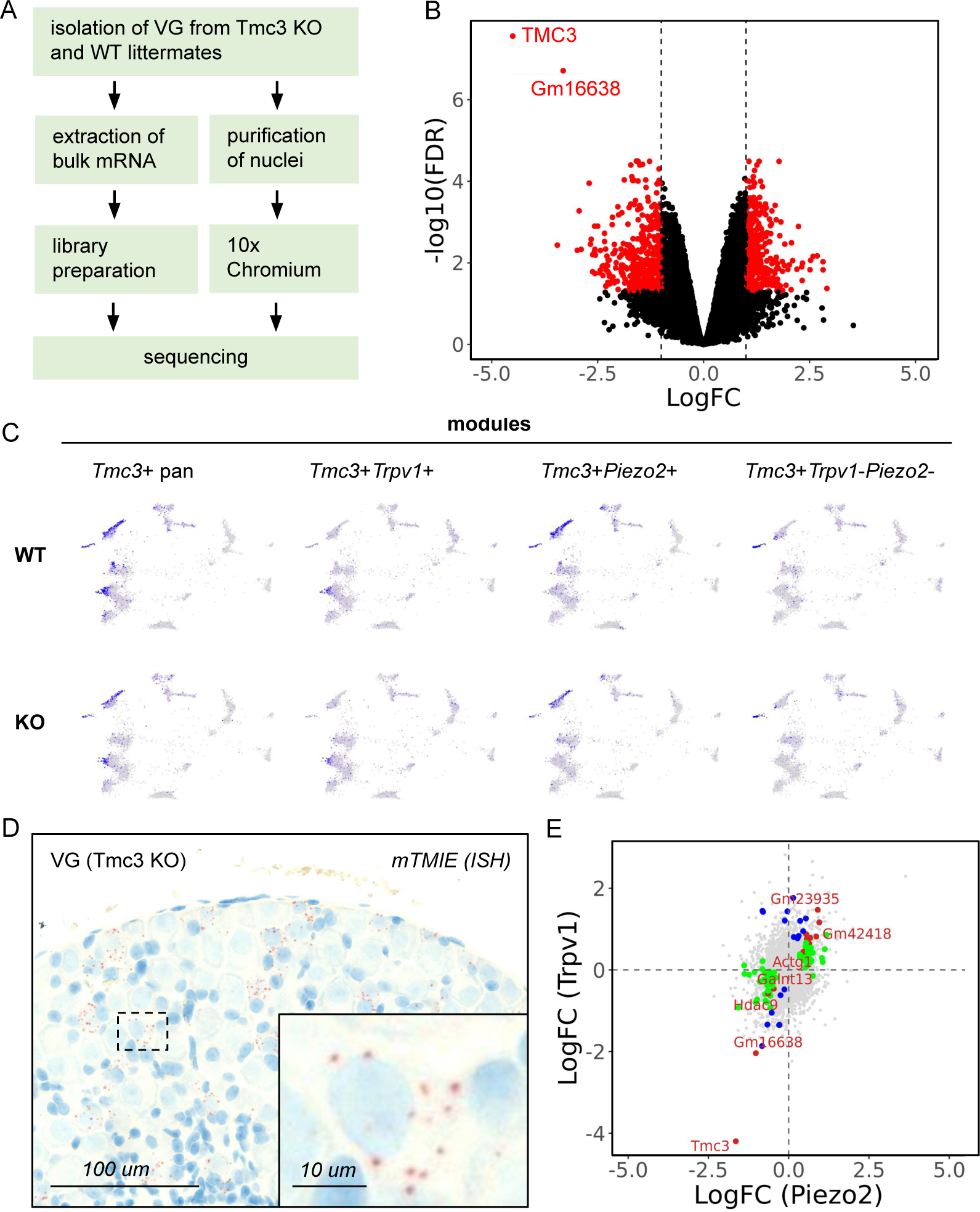
*Tmc3*+ deficiency reveals unique *Tmc3* dependent genes in the VG. (A) Schema of sample preparation. (B) Volcano plot from bulk-RNA seq analysis comparing wildtype and *Tmc3* KO nodose neurons. Genes that were significantly changed (FDR < 0.01 & LogFC > 1) are colored in red. Selected genes shown in plot. (C) UMAP representation of 21,311 single nuclei from VG projected onto coordinates from Fig 2. Blue color represents the level of specific *Tmc3* modules. (D) Detection of *Tmie* mRNA in *Tmc3* KO VG via ISH. Red color reflects positive *TmiE* signal, blue color reflects nuclei (E) 4-way plot of two separate differential expression analysis. x-axis shows log2 fold change from *Tmc3*-wt vs *Tmc3*-ko comparison of *Tmc3+Piezo2+* population; y-axis shows log2 fold change from *Tmc3* WT vs *Tmc3* KO comparison of *Tmc3+Trpv1+* population. Dots labeled in red are significantly different (FDR < 0.05 & fold change > 1.5) in both analyses. Blue and green represent significant changes in only *Trpv1*+ (blue) or *Piezo2*+ (green) analysis.

To provide additional insight into potential downstream targets of *Tmc3*, we performed single nuclei sequencing of VG from WT and *Tmc3* KO mice. After filtering low quality and non-neuronal nuclei, we were able to sequence 21,311 VG neuronal nuclei, capturing expression of about 2000 genes per nucleus. We mapped these nuclei to the coordinates of the previous integrated single-cell dataset and examined the expression of genes within the conserved modules described above (Fig. S4a). We found that the average expression of genes from each gene module was largely unaffected in *Tmc3* KO cells (Fig. 3C, and Supplementary Fig S4b). Furthermore, we verified that expression of *Tmie* (22), one of the most correlated genes with *Tmc3*, is still present in *Tmc3* KO mice (Fig. 3D). This was consistent in both bulk and single-nuclei datasets (Supplementary Fig S4c). Similar to results obtained *Tmc1, Tmc2* and *Tmie* (20–22), our data suggests that *Tmie* is not transcriptionally regulated by Tmc3, and that *Tmie* has the potential to serve as a *Tmc3*-proxy for VG neurons in *Tmc3* KO mice. Together, these data highlight that loss of *Tmc3* does not significantly alter the identity of the sensory neurons in which it is expressed.

We next performed differential expression analysis of individual *Tmc3*+ clusters isolated from wildtype and *Tmc3* KO mice using a pseudobulk approach (see Methods). Interestingly, this analysis revealed significant gene expression changes within specific single cell clusters (Fig.3E; Supplemental Table 3 and 4; FDR < 0.05 and > 1.5 fold change). Some genes were consistently altered across both *Tmc3+Piezo2+* and *Tmc3+Trpv1+* clusters (*Slc8a1, Nbea, Hdac9, and Plcxd3)*. However, we also identified a number of genes that were uniquely altered in *Tmc3+Trpv1+* and *Tmc3+Piezo2+* clusters. These genes include *Camk1d* and *Ahnak* in *Tmc3+Trpv1+* neurons, and *Kcnj3* and *Pde4d* in *Tmc3+Piezo2+* neurons. These data suggest that perturbing *Tmc3* may alter the expression of select ion channels, and the differentially expressed genes could reflect cell-specific downstream targets of *Tmc3*. These data will be useful in understanding the specific sensory functions of these sub-populations of *Tmc3*-expression sensory neurons.

### Chemogenetic modulation of *Tmc3*+ VG sensory neurons induces broncho-constriction or dilation with drastic impact on overall lung function

Next we sought to visualize and quantify the *in vivo* effects of *Tmc3*+ VG neuron inhibition or activation on lung function and bronchial integrity. Cre dependent Designer Receptor Exclusively Activated by Designer Drugs (DREADD) mice have been successfully utilized for chemogenetic neuronal control in a variety of *in vivo* studies, including efforts applying DREADD technology to understand breathing disorders or enable further insights into cardiopulmonary biology (31–35). This prompted us to generate inhibitory (hm4di, “DREADD-Gi”) and activating (hm3dq, “DREADD-Gq”) Tmc3-Cre-DREADD mice (Fig. 4A). These mice are designed to facilitate specific modulation (activation or inhibition) of *Tmc3*-expressing neurons via intraperitoneal injection of the designer drug clozapine-N-oxide (CNO).

**Figure 4.**
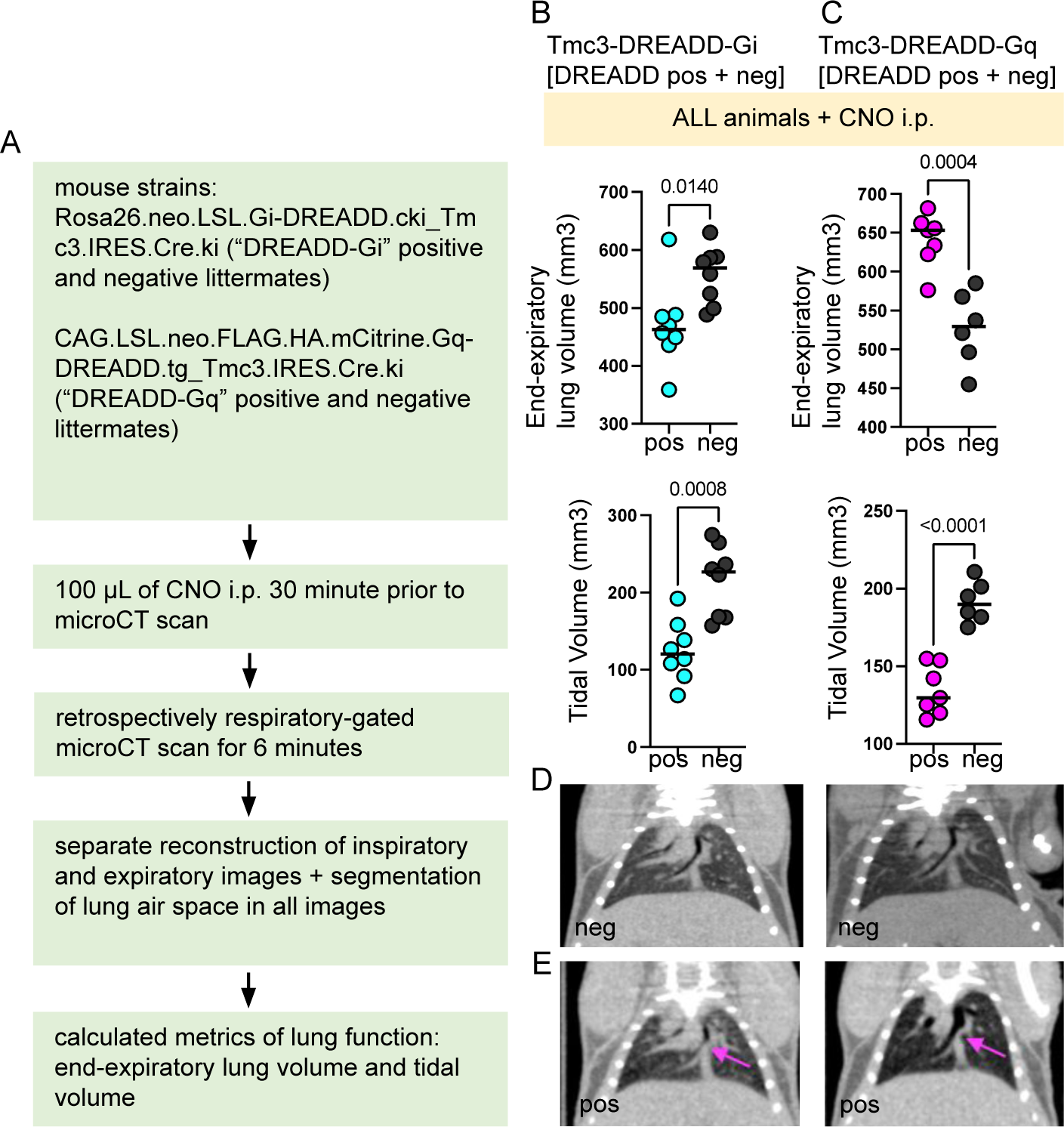
Chemogenetic modulation of *Tmc3*+ VG sensory neurons induces broncho-constriction or dilation with immediate consequences for the respiratory cycle. (A) Tmc3-DREADD-Gi/Gq mice or WT littermate controls were injected with CNO for subsequent microCT imaging to determine end-expiratory lung volume and lung density. (B-C) End-expiratory lung volume quantification across individual Tmc3-DREADD mice (n= 7-8) and WT controls (n= 6-8). (D) Airway constriction or dilation in comparison to WT control was visible at the bronchi-level (magenta arrow). (E-F) Both, activation (DREADD-Gq) and inhibition (DREADD-Gi) of *Tmc3*+ VG neurons led to a significant reduction in tidal volume compared to WT controls; individual values plotted with line representing median; P values are plotted on graph.

We combined the DREADD chemogenetics approach with micro computed tomography (microCT) imaging to visualize breathing patterns and overall lung integrity after inhibition or activation of *Tmc3*+ VG neurons. To rule out CNO-associated off target effects, all animals received CNO i.p. prior to downstream testing. This included DREADD-Gi and Gq positive mice (pos) as well as DREADD-Gi and Gq negative littermate controls (neg). All animals were lightly anesthetized to facilitate precise microCT imaging without affecting voluntary respiratory function. We excluded any respiratory measurements that would require involuntary or forced ventilation of the lungs (like inspiratory reserve or residual lung volume). Post CNO injection, Tmc3-Cre DREADD-Gi and Gq pos and neg mice were subjected to a retrospectively respiratory-gated microCT scan allowing for real time visualization of respiratory processes. These recordings were then used to calculate and quantify the specific lung performance parameters end-expiratory lung volume and lung tidal volume. A single respiratory cycle is characterized by completion of inhalation and exhalation. The end-expiratory lung volume is the amount of air remaining in the lungs at the end of a normal exhalation. The tidal volume is the exact amount of air that is exchanged (passing in and out of the lungs) during a single respiratory cycle.

CNO treatment of Tmc3-DREADD-Gi positive mice should cause inhibition of *Tmc3*+ VG neurons. Given the profound innervation of the lungs with *Tmc3*+ neurons we suspected their inhibition would have an immediate effect on lung function when assessed by gated microCT. Strikingly, intraperitoneal CNO injection into Tmc3-DREADD-Gi positive mice induced visible contraction of the lungs and an inability to expand the airways to their full capacity when compared with equally CNO-injected DREADD-Gi negative littermate controls (Fig. 4B). This state of locked contraction lasted for ∼1.5h and was accompanied by a significant decrease in end-expiratory lung volume compared to DREADD-Gi negative littermate controls (Fig. 4B). As expected, the decrease in lung volume of CNO-injected DREADD-Gi positive mice also affected their ability to exchange air during a respiratory cycle. This became evident by a significant reduced tidal volume of CNO-injected DREADD-Gi positive mice in comparison to equally treated DREADD-Gi negative littermate controls (Fig. 4B).

Conversely, intraperitoneal CNO injection should cause activation of *Tmc3*+ sensory neurons in Tmc3-DREADD-Gq positive mice but not in their Tmc3-DREADD-Gq negative littermates. CNO treatment of Tmc3-DREADD Gq positive mice induced dilation of lungs including bronchi for ∼ 1.5h visible by microCT (Fig. 4C). The ongoing airway dilation drastically interfered with the ability of Tmc3-DREADD-Gq mice to properly exhale air from their lungs. This resulted in air staying trapped in the lungs at the end of each exhalation visible by an elevated end-expiratory lung volume when compared to CNO treated DREADD-Gq negative controls (Fig. 4C). In addition, the lungs of CNO-treated Tmc3-DREADD-Gq positive mice remained at their maximum holding capacity, greatly interfering with inhalation of additional air. Ultimately, CNO-injected Tmc3-DREADD-Gq positive mice were not able to properly exchange air from the lungs. This inability to complete a full respiratory cycle caused a significant reduction in tidal volume when compared to CNO treated Tmc3-DREADD-Gq negative littermate controls (Fig. 4C). Notably, inhibition or activation of Tmc3+ VG neurons in Tmc3-DREADD-Gi and Gq positive mice induced significant airway narrowing or dilation, respectively, that was visible in individual bronchi when compared with DREADD negative littermate controls (Fig. 4D-E, pink arrow) - further emphasizing the impact of *Tmc3*+ neuron-dependent regulation of lung function. The effect of CNO on both the Tmc3-DREADD-Gi and Gq positive animals lasted for around 1.5h. Afterwards their breathing patterns returned to normal with no visible difference to their DREADD negative littermates. Taken together these chemogenetic studies showed that *Tmc3+* VG neurons play a critical role in regulating a healthy airflow cycle, likely through contraction or dilation of substantial portions of the lungs.

## Discussion

Sensory interoception of the lungs is thought to be largely accomplished by vagal afferents with a higher complexity of innervation compared to that seen in other organs (1, 4–6). Through a combination of imaging and labeling experiments, we showed that *Tmc3* is expressed in a specific subset of nodose neurons that innervate the distal airways. Leveraging internally generated bulk and single cell data, coupled with externally generated single cell data, we identified distinct *Tmc3*-expressing sub-populations of VG neurons that express *Trpv1* or *Piezo2*, respectively, which could reflect sensory neurons with different interoception roles in the airway. Lastly, we presented functional data showing that either activating or inactivating the neurons that express *Tmc3* can have a profound impact on lung respiratory function.

We and others have identified that *Tmc3* is highly expressed within the VG (36). Our study further extends previous work as we show that *Tmc3* expression is restricted to just the nodose ganglia, a subset of the entire VG. Our results are consistent with the model that nodose VG neurons innervate distal airways, whereas jugular afferents innervate the proximal airways (4, 37–41) as well as esophageal mucosa and myenteric ganglia (42). Retrograde labeling presented in our analysis highlights that only a subset of VG neurons innervate the airways, which is in line with data estimating that up to 20% of VG innervation projects to the lungs (1, 43). Interestingly, our data also indicate that a small number of *Tmc3*+ VG neurons did not innervate the airways, which suggests that these neurons could innervate other organs. In fact, RNAscope and ProjectionSeq were recently used to show that *Tmc3*+ VG neurons might also innervate other organs such as the stomach and pancreas (12).

Our integrative analysis of publicly available VG single-cell RNA-sequencing datasets from three separate labs identified robust *Tmc3* expressing subpopulations of neurons and their respective gene modules. These modules provide hints of the potential function of *Tmc3*+ VG neurons. For example, the expression of *Piezo2* and *Trpv1* point to roles of these sensory neurons in mechanosensation and nociception, respectively*. Piezo1* and *Piezo2* are low-threshold mechanoreceptors of the respiratory tract (44, 45) and have been studied for their role in airway processes. Optogenetic activation of vagal *Piezo2* leads to a state of acute apnea reminiscent of the Hering-Breuer inspiratory reflex (11). Similarly, we find that *Tmc3*+ VG neuron activation forces the lungs, and specifically the bronchi, into a dilated state. In contrast, inhibition of all *Tmc3*+ VG populations (including *Tmc3+Piezo2+* VG) resulted in a significant decrease in end-expiratory lung volume with a simultaneous reduction in tidal volume. This result differs from *Piezo2* KO mice, which have increased lung tidal volume with a simultaneous lack in response to airway stretch (11). The difference between lung phenotypes seen through *Piezo2* KO mice or inhibition of *Tmc3*+ VG neurons could be due to the extensive innervation of the lungs by *Tmc3+* VG and the additional involvement of *Tmc3+ TRPV1+* lung innervating neurons in smooth muscle contraction (46). Engineering new models that enable precise control of these neurons may provide even deeper insights into additional functions of *Tmc3*+ vagal afferents, and might also highlight additional subsets of lung-bound sensory neurons. Further exploring mechanisms underlying neuronal control of airway dilation and constriction could lead to therapeutic alternatives to existing corticosteroid treatment regimens for respiratory traits associated with acute bronchoconstriction in asthma, COPD and emphysema.

The transcriptional programs specific to *Tmc3*+ VG neuron subpopulations contained genes associated with sensorineural function, and were also found expressed in hair cells of the inner ear. *Tmc3* belongs to a family of 8 transmembrane channel-like receptors and the precise function of *Tmc3* in the VG sensory afferents complex is not known. *Tmc1* and *Tmc2* are involved in mechanotransduction processes in the inner ear (22, 27, 30, 47, 48) with *Tmc1* being a sensor protein for hearing and balance (28) and mutations in the *Tmc1* gene resulting in hearing loss (49–51). *Tmie* and *Lhflp3*, genes that are expressed in inner ear cells and have been reported in the context of hearing loss (20–22), are also co-expressed with *Tmc3* in the nodose ganglia. This raises an interesting possibility that molecular mechanisms that require TMC family members could be utilized in different biological contexts, and could inform our understanding of the processes being regulated in distinct cell populations. Interestingly, recent work highlights that TMIE and TMC1/2 physically interact and are subunits of the mechanotransduction channel that functions in inner ear cells (20). A similar relationship could exist between TMIE and TMC3 in the nodose neurons, suggesting a role for TMC3 in sensing mechanical force in the airways.

Importantly, although our data were generated in mice, the findings from our study could inform human biology. Murine VG are composed of jugular and nodose ganglion fused together into a single complex, while these ganglia are anatomically separated in humans (1, 4, 52). The human nodose ganglion alone is considerably larger than the mouse vagal ganglion including a proportionally higher number of neuronal cell bodies (2, 53, 54). However, we see a similar distinct *TMC3* mRNA expression pattern in sections generated from human donor VG (Fig. S1B), suggesting the possibility that we can infer aspects of human airway interoception from our work on mouse *Tmc3*+ VG afferents.

In summary, our work integrates sophisticated single cell analysis approaches with complex imaging techniques to characterize specific sub-populations of sensory neurons that innervate the airways. Importantly, we provide functional data revealing that activation or inhibition of these neurons significantly modulates airway function. The results presented here lay out critical insights into the role that sensory neurons play in broncho-constriction and dilation, and provide novel mechanistic and therapeutic hypotheses that could be tested for treatment of airway disorders.

## Materials and Methods

### Animals

All experimental procedures involving animals were approved by Genentech’s Institutional Animal Care and Use Committee and adhered to the National Institutes of Health (NIH) Guide for the Care and Use of Laboratory Animals. Male or female *Tmc3*-Cre mice and their WT littermates were used at an age of 8-12 weeks. Additional C57BL/6N mice were acquired from Charles River Laboratories. The two DREADD mouse alleles were licensed from University of North Carolina, Chapel Hill, and also obtained from the Jackson laboratory: The *Rosa26*-lox-stop-lox-Gi-DREADD knock-in allele from JAX strain # 026219 (B6.129-Gt(ROSA)26Sor tm1(CAG-CHRM4*,-mCitrine)Ute /J) and the CAG-LSL-Gq-DREADD transgenic allele from JAX strain #026220 (CAG-LSL-Gq-DREADD B6N;129-Tg(CAG-CHRM3*,-mCitrine)1Ute/J).

The Tmc3.IRES.Cre.ki (see below for generation details) is under the control of the Tmc3 promoter and will express Cre in the *Tmc3*+ VG and all other tissues where *Tmc3* is potentially expressed. Both DREADD mouse strains were crossed with Tmc3.IRES.Cre.ki mice. This resulted in the following strains: CAG.LSL.neo.FLAG.HA.mCitrine.Gq-DREADD.tg_Tmc3.IRES.Cre.ki and Rosa26.neo.LSL.Gi-DREADD.cki_Tmc3.IRES.Cre.ki. The Rosa26-lox-stop-lox-tdTomato allele was licensed from the Allen Institute for Brain Science and imported from the Jackson laboratory (#007909, B6.Cg-Gt(ROSA)26Sor tm9(CAG-tdTomato)Hze /J) and crossed with Vglut2-ires-cre knock-in allele mice licensed from Harvard Medical School and obtained from JAX (#028863, B6J.129S6(FVB)-Slc17a6_tm2(cre)Lowl/MwarJ) resulting in Rosa26.LSL.tdTomato.cki_SLC17a6.IRES.Cre.ki mice. The *Tmc3* Knock-out/LacZ knock-in allele (C57BL/6N-*A^tm1Brd^ Tmc3^tm2a(KOMP)Wtsi^*/Mmucd; here called TMC3 KO) was imported from the Mutant Mouse Resource & Research Centers (MMRRC) at University of California, Davis (https://www.mmrrc.org/catalog/sds.php?mmrrc_id=50081) and maintained on a C57BL/6N background.

### Generation of Tmc3-Cre mice

The construct for targeting the Tmc3 locus in ES cells to generate the IRES-Cre allele was made using a combination of recombineering, gene synthesis and standard molecular cloning techniques. The resulting targeting vector enabled insertion of an IRES-Cre FRT-*Pgk1*-Neo-FRT cassette after the Tmc3 stop codon at genomic position mm10 chr7:83,622,947. The vector was confirmed by DNA sequencing, linearized with NotI and used to target C57BL/6 C2 ES cells using standard methods (G418 positive and gancyclovir negative selection). Positive clones were identified using PCR and taqman analysis, and confirmed by sequencing of the modified locus. Correctly targeted ES cells were transfected with a Flpe plasmid to remove the Neo selection marker and create the final Tmc3 IRES-Cre allele. Validated ES cells were injected into blastocysts using standard techniques, and germline transmission was obtained after crossing resulting chimeras with C57BL/6N females.

### Passive Intra-tracheal Inhalation (PITI)

Mice were anesthetized in an appropriately sized induction chamber with 5% sevoflurane and 1.5-3 L O2/min. Upon a confirmed deep plane of anesthesia each mouse was placed on its back, on a mouse intubation stand (Kent Scientific, ETI-MSE-01) and suspended by its top front teeth on a small loop of string. A small cotton swab was used to gently roll the tongue out of the mouth and to the side, where it was held securely. A smooth trimmed pipette tip (the “introducer”) was inserted into the animal’s mouth, with the narrow end of the introducer positioned over the tracheal opening. The i.t. treatment was pipetted into the middle of the introducer, where it remained due to surface tension. The liquid only exited the introducer and entered the tracheal opening upon inhalation with the animal’s own breath. The mice were removed from the stand and recovered on a 37 °C heatpad, and monitored until fully conscious and ambulatory. NIRF imaging confirmed even distribution of PITI administered dye in the lungs (Fig. S5).

### Confirmation of Tmc3 antibody specificity in transfected U2-OS cells

U2-OS cells were grown and maintained in DMEM:F12, 10% FBS, 2mM Glutamine media according to ATCC recommendations and seeded at 150k cells/well in chamber coverslips. Transfections were done using Lipofectamine 2000 (Thermo Fisher Scientific) and 500 ng final concentration of pRK5-mTmc3-FLAG constructs or empty vehicle according to manufacturer’s instructional and fixed 48 hours later in 4% PFA in PBS. Monoclonal anti-FLAG M2 antibody (F3165, Sigma) was used at 1:1000, rb-anti-Tmc3 (HPA040265, ATLAS) was used at 1:200, Phalloidin-657 stain (Abcam) was used at 1:1000 and secondary F(ab’)2 Fragment antibodies (dk-anti-rb-488, 711-546-152; dk-anti-rb-647, 711-606-152; dk-anti-m-488, 715-546-151; dk-anti-m-647, 715-606-151; Jackson ImmunoResearch) were used at 1:500. DAPI was added at 1:1000 and cells were mounted with ProlongGold glass antifade medium at room temperature overnight and analyzed using a Leica SPE confocal microscope.

### Preparation of Vagal Ganglia (VG) cell suspension for IF

Mouse VG were harvested and pooled from 5 mice per treatment group. Nodose homogenate was obtained by carefully pulling whole VG from three mice. Mice were sacrificed in the morning by carbon dioxide inhalation followed by decapitation. After the ganglia were removed, the isolated ganglia were immediately transferred into a 24-well plate containing digestion solution (1 ml high glucose DMEM and 100 μl Collagenase Type IV (Sigma; C1889-50 mg) and incubated at 37°C for 40 min. The nodose ganglia were then transferred to a well containing 1 ml HBSS- and 100 µl Trypsin (Sigma; T9935-50 mg) and were incubated 5 min at 37°C. After digestion, the nodose ganglia were gently washed three times with Hibernate A medium (Gibco Cell Therapy Systems, A13705-01), preventing any tissue breakdown. The following trituration step was performed in 500 μl of Hibernate A by using a 1-ml pipette (pipette up and down 10–15 times) followed by a 200-μl pipette (pipette up and down 15–20 times). This procedure generated a cloudy solution containing dissociated cell suspension. For downstream IF imaging and quantification 900 µl Hibernate A was slowly added to the side of the chamber coverslip wells and cells were recovered O/N at 37° C before being fixed with 2% PFA in culture medium (prewarmed to 37° C) for 15 minutes. Chamber slides were then removed from incubator and culture medium was replaced with 4% fixative for another 10 min at room temperature. Cells were then washed with DPBS containing 50 mM glycine or NH_4_Cl for at least 5 changes over the course of 15 min. Blocking solution (5% BSA, 0.5% gelatin, 0.05% Tween 20 in DPBS; preferably filtered) was added for 15 min at RT before primary rb-anti-TMC3 (ATLAS HPA040265) and ms-anti-Beta3TUB (ABCAM, ab7751) antibody was added at 1:200 and VG cells were incubated at 4° C O/N on a lightly tilting tissue culture platform. The next day cells were washed with blocking solution with a minimum of 3 washes for a total of 30 min. Cells were then incubated with secondary antibodies (488 goat anti-rabbit F(ab’)2 fragment, 111-546-003 and 594 donkey anti-mouse F(ab’)2 fragment, 715-586-150; Jackson ImmunoResearch) diluted 1:500 in blocking solution for 30 minutes at room temperature. After another washing series with blocking buffer (30 minutes, at least 5 washes) and DPBS (15 minutes, at least 3 washes) slides were mounted with ProlongGold glass antifade medium at room temperature overnight and analyzed using a Leica SPE confocal microscope. VG neurons from 5 Fast Blue treated (PITI) WT mice were pooled and evenly plated on 3 coverslips. Individual b3tub+ cells were counted and assessed for FB and Tmc3 signal. For the retrograde tracing of lung-innervating VG neurons mice received a volume of 40 µl Fast Blue solution (350 µM and 1% DMSO in 0.9% saline, 73819-41-7, Polyscience Inc) or 40 µl DiD solution (0.5 m/ml in 0.9% saline, D7757, Thermo Fisher Scientific) via PITI on two consecutive days. For FACS Calcein-AM (PK-CA707-80011-3, Promokine) was added 1:1000 right before the analysis.

### Retro-orbital sinus injection of AAV reporter for *in vivo* fluorescence labeling purposes

Mice were anesthetized in an appropriately sized induction chamber with 5% isoflurane and 1.5-3 L O2/min. Once visually anesthetized, the mouse was transferred to a nose cone with 2.5% isoflurane. The mouse was placed in lateral recumbency and a drop of 1% proparacaine was instilled in the eye to be injected. Upon confirmation of a deep plane of anesthesia, indicated by the absence of toe pinch reflexes, gentle pressure was applied to the eye socket to partially protrude the eyeball. A 30-gauge needle was carefully introduced, bevel down, at an angle of approximately 45° in the retro-orbital sinus. AAVphp.s-CAG-DIO-dTOM (36 ul, 2.8 x 10^13 GC/ml) was injected. This inoculum was aqueous, not viscous. If blood was observed after injection, manual pressure with a sterile gauze was applied to ensure hemostasis. A drop of ophthalmic lubricant was instilled in the eye. The mouse was monitored until completely recovered from anesthesia and fully ambulatory.

Construct information: https://en.vectorbuilder.com/vector/VB200806-1011nvm.html AAV serotype is php.s reference: https://pubmed.ncbi.nlm.nih.gov/28671695/

### Processing and imaging of whole-mount tissue

Processing and imaging of whole-mount tissue was done as previously described (Balestrini et al. 2020). Briefly, animals were anesthetized using isoflurane and perfused with PBS/Heparin (5 U/ml) followed by tissue fixation using paraformaldehyde 4% (55). Lung tissue was harvested and subjected to immunolabeling using primary conjugated antibodies and a previously published staining protocol avoiding methanol treatment steps (56). Tissue clearing was performed using the FluoClearBABB approach (57), and whole-mount images were then acquired using a Leica SP8 microscope equipped with a white light laser and a Leica BABB immersion lens (HCX APO L 20×/0.95 immersion media). Acquired data were visualized on a power workstation using Imaris (Bitplane). Vagus nerve, VG and DRG soma were harvested, fixed in 2% PFA-PBS for 10 minutes at 37° C and analyzed for reporter expression by whole-mount IF (40x, oil) using a Leica SPE confocal.

### Lung function analysis of Tmc3-DREADD mice using in vivo microCT

Clozapine N-oxide (CNO) (Fisher Scientific, A3317-50) solution was prepared at a final concentration of 700 µg/ml CNO in 0.9% saline, 0.5% DMSO. All mice (n=7-8 per DREADD genotype and n=6 control animals) were injected with 100 µL of CNO i.p. 30 minute prior to microCT scan. All animals received a retrospectively respiratory-gated scan for 6 minutes. Inspiratory and expiratory images were reconstructed separately and lung air space was segmented in all images. The following metrics of lung function were measured: tidal volume (TV), normalized tidal volume (NTV, 100%*TV/EEV) and end-expiratory lung volume (EEV). Anesthesia (2-3% sevoflurane) was adjusted to maintain respiration rate in ideal range (∼60-140 breath per minute). Body temperature was maintained at 37° C by use of a warming pad.

### *In situ* hybridization

Vagal ganglia from mice or donors were obtained as described above. Nerve endings and connective tissues were trimmed and ganglia were placed either in OCT molds and frozen rapidly or processed as FFPE tissues. Human vagal ganglia were acquired from commercial sources under warranty that appropriate Institutional Review Board approval and informed consent were obtained. 5-10 micron sections were cut using a cryostat and placed on slides for downstream RNA labeling studies. RNAScope 2.5 LS Duplex Kit or Multiplex Fluorescent Kit v2 (Advanced Cell Diagnostics) was used per manufacturer’s recommendations for fresh-frozen or FFPE samples with the following alterations for fresh frozen samples: during pretreatment, sections were treated with hydrogen peroxide for 10 min and Protease IV for 2 0min prior to the addition of relevant probes. Opal dyes 690, 570, and 520 (Akoya Biosciences) were used for fluorescence and after the final HRP block step, samples were stained with DAPI solution for 1min. followed by mounting with ProLong Gold Antifade Mounting Solution (Thermo Fisher Scientific, Cat#P36961). Probes used for RNAscope 2.5 LS Duplex Kit include Mm-TMC3 (Cat#426308) and Hs-TMC3 (#809228) and probes for RNAscope Multiplex Fluorescent Kit (ACD) include Mm-Gal (Cat#400961), Mm-Tubb3 (Cat#423391) and Mm-TMC3 (Cat#426301).

### Low input bulk RNAseq of VG and DRG

Low input bulk RNAseq was performed using VG and thoracic DRG from individual animals (n=5). VG and DRG were transferred into an 1.5ml tube containing one Tungsten Carbide bead (3mm, Qiagen, #69997) and 1 mL of Qiazol lysis reagent (Qiagen, #79306) and homogenized for 2 x 3 minutes, at a frequency of 28/s @ 4°C (Tissue Lyser II, Qiagen). After this step the samples were adjusted to RT for 5 minutes and processed using the RNeasy Lipid Tissue Mini Kit (Qiagen) as described in the user manual. After elution of the RNA in 12 μl RNase free water, the RNA concentration was determined via Nanodrop and up to 100 ng was used as an input material for library preparation using the Clontech Ultra Low kit (Clontech).

### Bulk RNA-Seq alignment

The fastq sequence files for all RNA-seq samples were filtered for read quality and ribosomal RNA contamination. The remaining reads were then aligned to the mouse reference genome (GRCm38) using the GSNAP alignment tool (Wu and Nacu, 2010). Alignments were produced using the following GSNAP parameters: ‘-M 2 n 10 -B 2 -i 1 N 1 w 200000 -E 1 –pairmax-rna = 200000 –clip-overlap’. These steps, and the downstream processing of the resulting alignments to obtain read counts and normalized Reads Per Kilobase Million (nRPKMs) per gene (over all exons of RefSeq gene models), are implemented in the Bioconductor package, HTSeqGenie (v 3.12.0) (Pau and Reeder, 2014). Only uniquely mapped reads were used for further analysis.

### Bulk RNA-seq differential gene expression

Read counts were adjusted based on DESeq2 estimated sizeFactor (Love et al., 2014) and weighted by their mean-variance relationship as estimated by voom. Differential gene expression was performed using the limma empirical Bayes analysis pipeline described in the R package limma (Ritchie et al., 2015). Significant differences were defined as genes with p-value adjusted less than 0.01 and fold change greater than 2. p-values were adjusted using the false discovery rate (FDR) method following the Benjamini-Hochberg procedure.

### Bulk RNA-seq data presentation

For heatmaps, genes were log2-transformed, then scaled and zero centered using the z-transform. An arbitrary floor of −4 was set for genes with a log2 RPKM value of less than −4. Heatmaps were plotted using the superheat package in R.

### Single nuclei isolation

Due to the low number of nodose neurons we pooled VG from 3 mice for nuclei isolation per analyzed sample (4 samples total). Harvested VG were incubated in 0.5 mL Lysis buffer (10 mM NaCl; 10 mM Tris pH 7.5; 3 mM MgCl2; 0.2 U/mL RNase inhibitor) for 5 min on ice. VG were then carefully homogenized using a dounce homogenizer and spun at 300 g for 5 minutes. Supernatant was discarded and the pellet was resuspended in 0.5 mL PBS+ 3% BSA+ 0.2 U/mL RNase inhibitor containing 5 uL RNase free DNaseI and incubated at RT for 2 minutes. The solution containing nuclei was gently triturated, pelleted and resuspended in PBS+ 3% BSA+ 0.2 U/mL RNase inhibitor. Isolated nuclei were injected into Chromium for GEM generation and subsequent library construction (V3 of the Chromium 3’ EXP reagent kit; 10x Genomics). cDNA of the first RT reaction as well as the final library was QC’ed via High Sensitivity DNA chip bioanalyzer (Agilent).

### Single cell data curation and processing

When possible, the rawest form of the data was curated from publicly available for datasets. For Kupari et al., processed bam files were converted back to fastq. Prescott et al raw fastq were downloaded. All fastq files were aligned to the mouse reference transcriptome (GRCm38) via the 10x software Cellranger (v3.1.0) using default parameters to generate the raw feature and barcodes matrices. For Zhao et al., we directly downloaded the author provided raw feature and barcode matrix output from 10x for each individual sample.

### Single cell data integration and cluster analysis

For each individual dataset, raw 10x output was imported into R for downstream analysis. We used an outlier based approach to identify low-quality cells defined as those with: 1) Low total counts, indicating that library preparation or sequencing depth was suboptimal; 2) Low numbers of detected features, a slightly different flavor of the above reasoning; 3) High proportions of counts in the mitochondrial (or spike-in) subsets, representing cell damage. Outliers are defined on each metric by counting the number of median absolute deviations from the median value across all cells. This assumes that most cells in the experiment are of high (or at least acceptable) quality. For the total counts and number of detected features, the outliers are defined after log-transformation of the metrics. This improves resolution at low values and ensures that the defined threshold is not negative. Outlier identification is done separately in each batch. After QC filtering, we re-use the total sum for each cell to compute size factors based on the library size. We divide each cell’s counts by its size factors and then log-transform. This is done after blocking on the batch, where scaling is performed to match the coverage of the lowest-coverage batch. We fit a mean-dependent trend to the variances of the log-expression values. This is done in each batch and the average statistic across batches is reported for each gene. We then define the highly variable genes as those with the largest residuals. We run a PCA to compress and denoise the dataset by only extracting the top few PCs. This is weighted to ensure that both small and large batches have equal opportunity to contribute to the rotation vectors. The aim is to ensure that variations unique to small batches are still properly represented. PCs were used in place of the per-gene log-expression matrix in downstream (distance-based) calculations. Distances in PC space can be used as proxies for those same distances in the original expression space.

For each individual dataset, we treat each sample as a batch and perform batch correction on the PCs using a mutual nearest-neighbors approach. This identifies matching populations across batches and merges them together while keeping unique (batch-specific) populations separate. The output is a set of corrected PCs that we will use in place of the original PCs in our downstream analyses.

We next used an automated annotation tool, SingleR, to roughly annotate the cell types. Clusters with >90% neuronal identity were retained for downstream integrative analysis. We leveraged the “celldex” R package, which provides a collection of reference expression datasets with curated cell type labels.

After each individual dataset has been batched-corrected, merged, and filtered for neurons, we used the Seurat package to integrate between the 3 datasets using the FindIntegrationAnchors() and IntegrateData() functions with dims=1:30 and anchor.features=2000.

To verify the reliability of these clusters, we performed an independent computation approach, MetaNeighbor (18), to quantify cell type replicability across datasets. We used the MetaNeighbor package (v1.20.0) with default parameters and set fast_version = TRUE.

### Single cell gene network analysis

We used the Hotspot software (v1.1.1) to identify unique modules across each dataset. For each dataset, hs.create_modules() function was used to generate unique modules, but different gene thresholds were used to achieve similar module counts across datasets. For Kupari et al, we used min_gene_threshold=80; Prescott et al. min_gene_threshold=80; Zhao et al min_gene_threshold=80. Module scores were calculated using the hs.calculate_module_scores() function with default options, and plotted to determine the localization with Tmc3. Genes that were present in at least 2 separate modules across datasets were retained.

### Single nuclei data processing

Raw sequencing reads were processed and aligned to the mouse reference transcriptome (GRCm38) via the 10x software Cellranger (v3.1.0) using default parameters to generate the raw feature and barcodes matrices. CellBender (v0.2.0) was used to remove ambient RNA effects from raw gene-by-cell count matrices for each sample using the remove-background function with the following parameters: epochs=200, fpr=0.01, total-droplets-included=30000, and expected-cells=number of cells called by CellRanger. CellBender produces processed and filtered gene-by-cell count matrices corrected for ambient RNA.

Individual samples were merged and analyzed using Seurat (v3.1.1) following the standard workflow. Briefly, cells with less than 500 UMIs or mitochondria RNA content greater than 25% were filtered. For each cell, the feature expression counts were normalized by the total expression and log transformed. Next, feature selection was determined by computing the top 1000 highly variable genes using the vst method. For single-cell data, 2000 highly variable genes were used. Finally, count values per gene was zero center scaled prior to dimension reduction analysis.

### Single nuclei cluster analysis

Principal component analysis was performed on scaled data and the top 30 components were used for UMAP analysis. Clusters were identified by first constructing a KNN graph based on the euclidean distance in of the top 30 components in PCA space, then using a Louvain algorithm to iteratively group cells together. A range of resolution parameters were used in the FindClusters function and the best resolution was visually determined based on marker expression and known biology.

### Single nuclei pseudobulk analysis

Individual pseudobulk samples were generated for each animal by aggregating counts across all cells using the aggregateAcrossCells() function in the scater package (v1.24). We also generated pseudobulk samples for each animal and each cluster by aggregating counts across all cells for each cluster. Differential expression analysis was performed on pseudobulk samples using the voom-limma method.

### Single nuclei cell annotation

Individual clusters were initially labeled using the automated annotation tool, SingleR using the same “celldex” reference as single cell analysis. We further cross-referenced the cluster labels with previous datasets (e.g. Kupari et al., Prescott et al.). We downloaded processed data from the Kupari dataset and used Seurat’s LabelTransfer function to superimpose previously defined cluster subtypes onto our nuclei data.

### Neuropeptide correlation

To search for neuropeptides associated with Tmc3 expression, we performed Spearman correlation between Tmc3 expression and expression of a list of curated neuropeptides.

### Accession number

RNA-sequencing and single nuclei RNA-seq data has been deposited to the Gene Expression Omnibus (GEO) under accessions GSEXXXXXX and GSEXXXXXX, respectively.

### Statistical analysis

Wetlab experimental data were analyzed using GraphPad Prism 9. For the microCT lung function experiments individual dots on each graph represent individual animals and not technical replicates. snSeq data was corroborated by analysis of 2 independent published data sets. Unpaired t test was used for the mouse data shown. A P value less than 0.05 was considered significant.

## Acknowledgements

We thank members of the Immunology department for discussions; Jessica Montealvo for procurement and management of FFPE human nodose ganglia tissue; Jian Jiang and Charles Jones III for histologic processing and sectioning of FFPE tissues; Soren Warming and Matthew Ho for assistance with mouse strain management; Meron Roose-Girma and team for generation of the TMC3-Cre mouse strain; Alvin Gogineni, Martin Weber and Eric Suto for additional experimental support; facility staff at Genentech for vivarium maintenance and core facility assistance. This work was supported by Genentech.

## Supplementary Figure Legends

**Figure S1.**
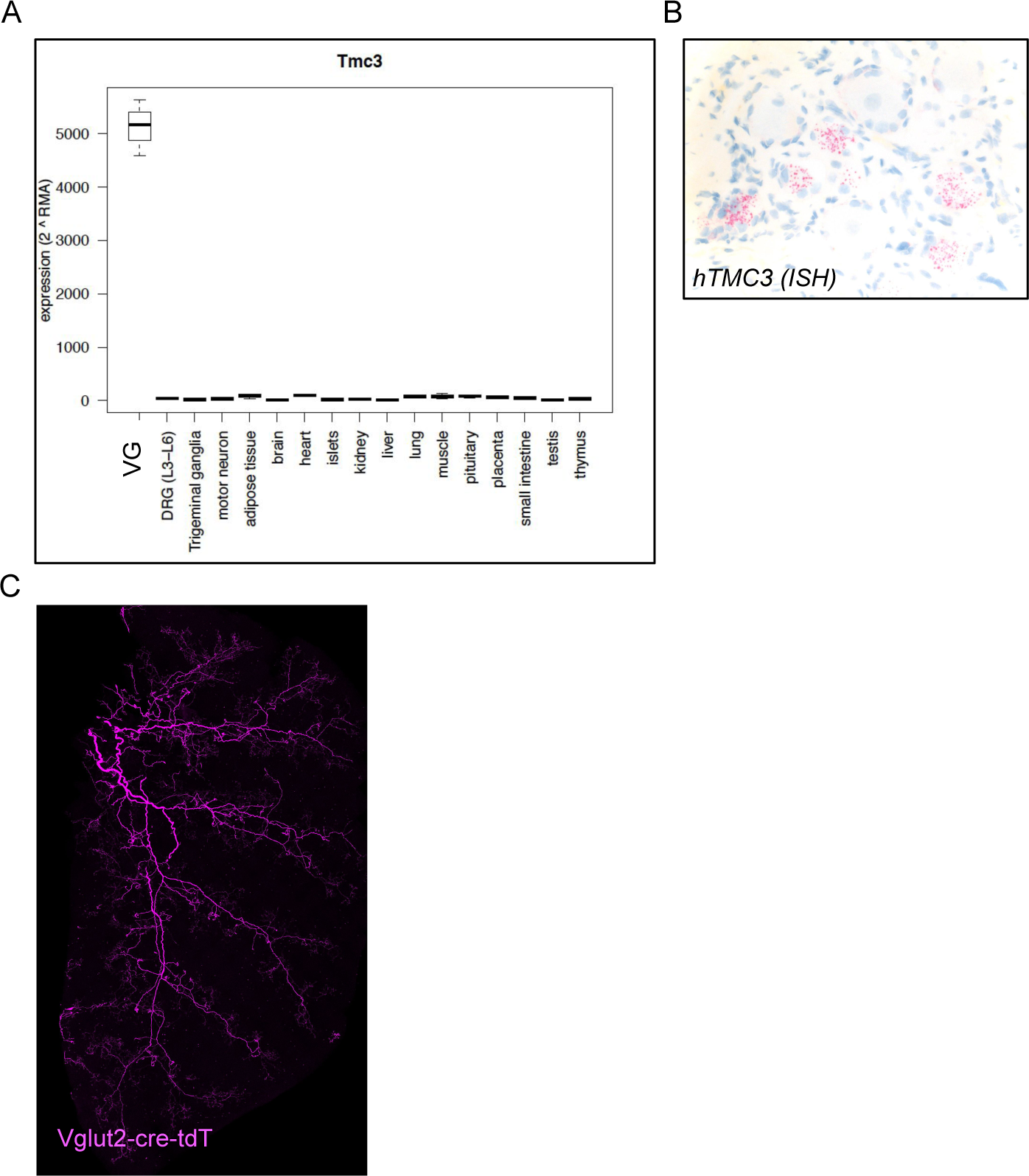
(A) *Tmc3* gene expression by qRT-PCR in various tissues, as indicated. (B) Representative ISH indicating *Tmc3* transcripts in sections of human nodose ganglia tissue. (C) Lung-Innervation was assessed by 3D imaging of cleared whole lungs from Vglut2cre-tdTomato mice.

**Figure S2.**
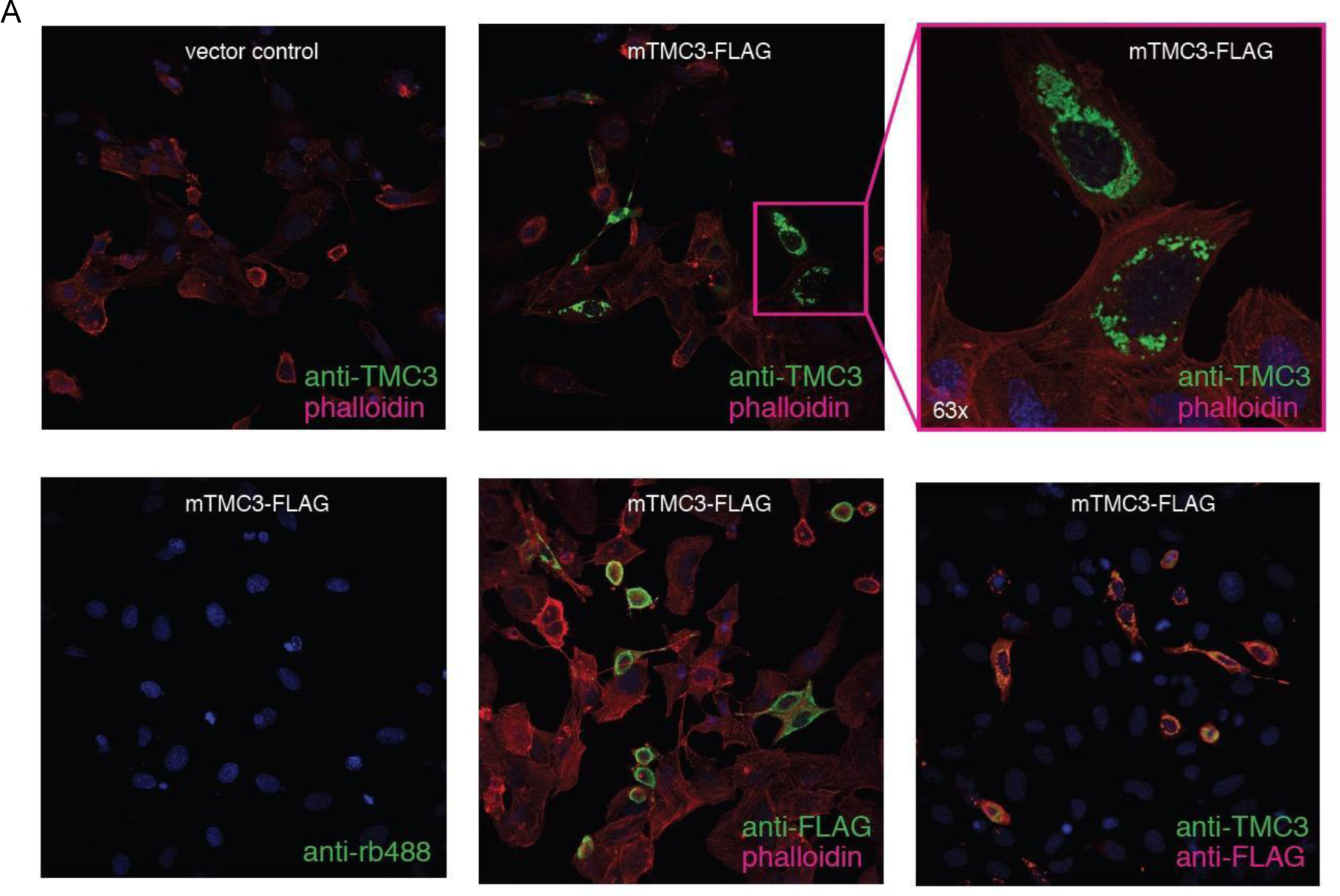
(A) Tmc3 antibody specificity was confirmed in U2-OS cells transfected with Tmc3-FLAG constructs or vehicle control and stained with anti-Tmc3 antibody or/and anti-FLAG antibody. Nuclei were visualized via DAPI counterstain; phalloidin staining as indicated in individual tiles.

**Figure S3.**
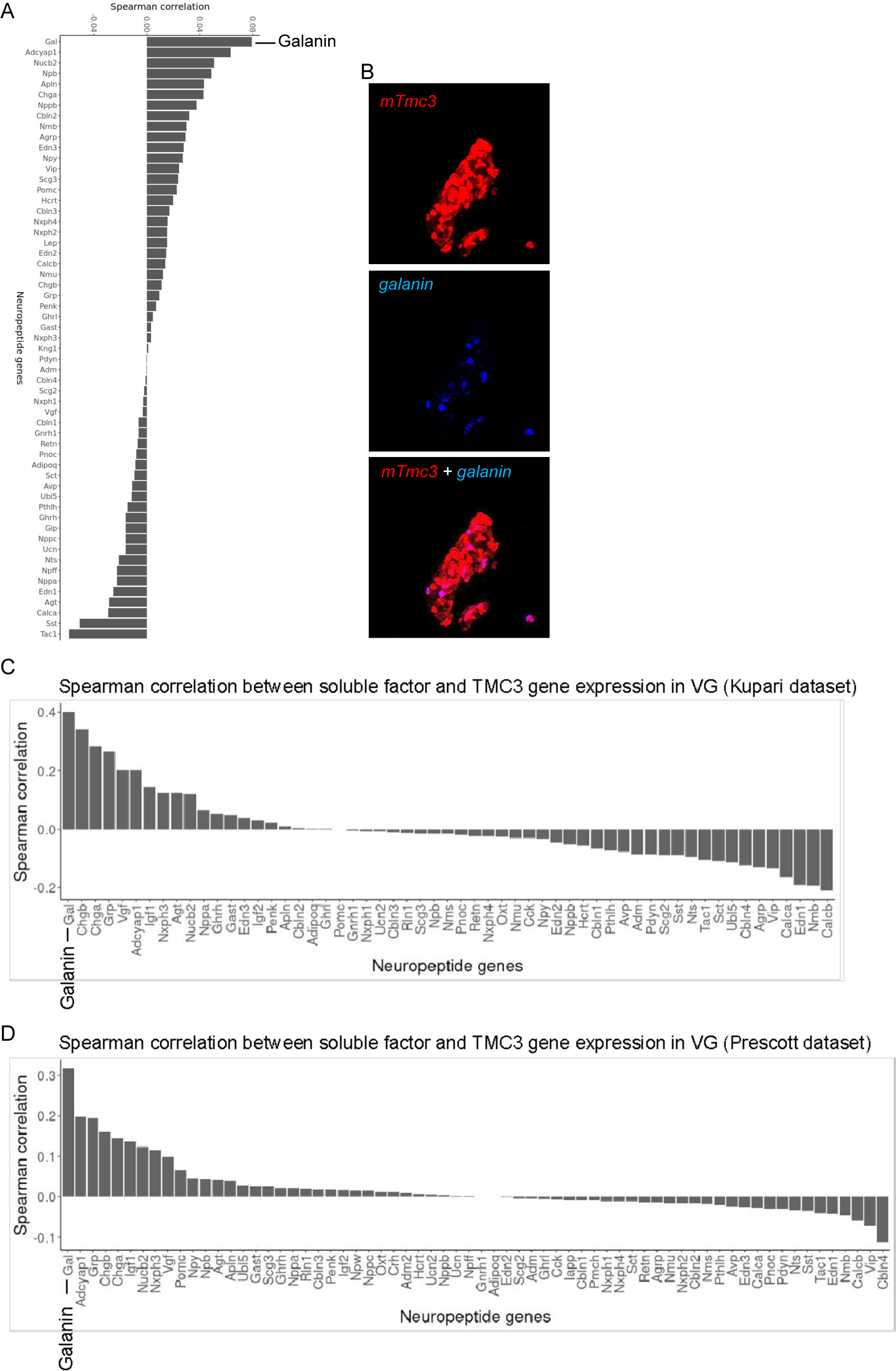
(A) Spearman correlation between soluble factors/neuropeptide genes and *Tmc3* gene expression in VG using snSeq dataset. (B) Co-expression of *Galanin* and *Tmc3* mRNA in VG soma identified via ISH. (C-D) Spearman correlation between soluble factors/neuropeptide genes and *Tmc3* gene expression in published scSeq datasets from Kupari et al. and Prescott et al. (as indicated).

**Figure S4.**
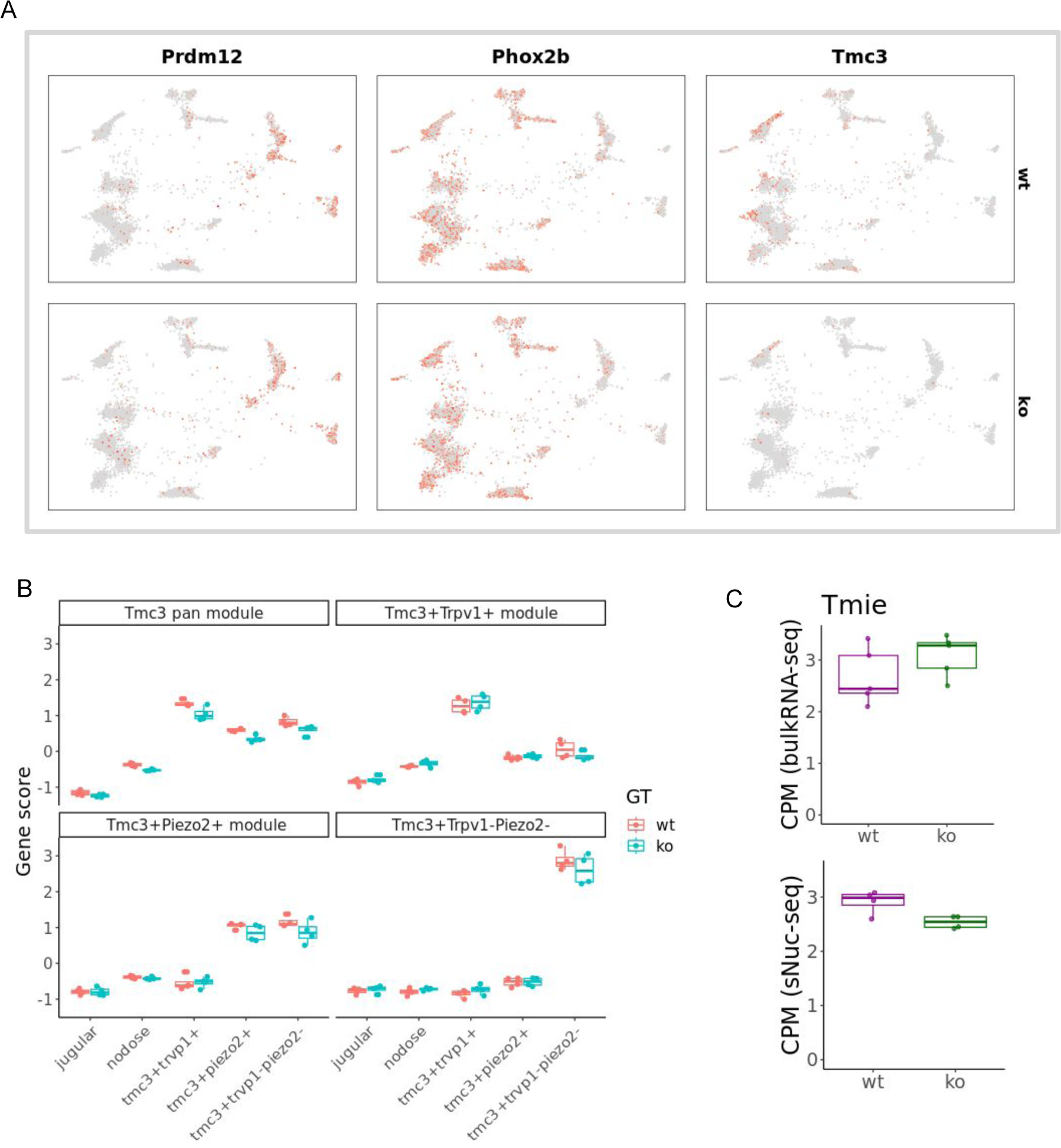
(A) UMAP representation of the sNuc-seq dataset mapped onto the coordinates of the integrated data in Figure 2. Relative expression of *Prdm12*+ (jugular), *Phox2b*+ (nodose), and *Tmc3*+ neurons is shown by the intensity of color. (B) Boxplot showing the gene score of different *Tmc3* modules in different animals summarized as pseudobulk samples. x-axis refers to different classes of VG neurons. (C) Boxplot of *Tmie* expression in bulk (top) and sNuc-seq pseudobulk samples between WT and KO.

**Figure S5.**
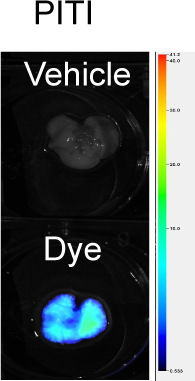
NIRF imaging of whole isolated lungs after PITI administration of vehicle (PBS) or dye as indicated.

